# Specific attributes of the V_L_ domain influence both the structure and structural variability of CDR-H3 through steric effects

**DOI:** 10.1101/2023.05.16.540974

**Authors:** Bora Guloglu, Charlotte M. Deane

## Abstract

Antibodies, through their ability to target virtually any epitope, play a key role in driving the adaptive immune response in jawed vertebrates. The binding domains of standard antibodies are their variable light (V_L_) and heavy (V_H_) domains, both of which present analogous complementarity-determining region (CDR) loops. It has long been known that the V_H_ CDRs contribute more heavily to the antigen-binding surface (paratope), with the CDR-H3 loop providing a major modality for the generation of diverse paratopes. Here, we provide evidence for an additional role of the V_L_ domain as a modulator of CDR-H3 structure, using a diverse set of antibody crystal structures and a large set of molecular dynamics simulations. We show that specific attributes of the V_L_ domain such as CDR canonical forms and genes can influence the structural diversity of the CDR-H3 loop, and provide a physical model for how this effect occurs through inter-loop contacts and packing of CDRs against each other. Our study provides insights into the interdependent nature of CDR conformations, an understanding of which is important for the rational antibody design process.

## Introduction

A key part of the adaptive immune response of jawed vertebrates is the use of antibodies to recognise and bind to extracellular antigens, thereby driving part of the downstream specific immune response. Standard antibodies are composed of two light and two heavy chains, forming a heterotetrameric complex, with two identical F_V_ regions, which contain the antigen-binding site. This region is made up of the variable heavy (V_H_) and light (V_L_) domains, each containing three loops referred to as complementarity-determining regions (CDR1-3) that make up the majority of the antigen-binding residues called the paratope (1). The remainder of the variable domains of both V_H_ and V_L_ is referred to as the framework region. Through two major processes, namely V(D)J recombination and affinity maturation, the structure of the paratope is fine-tuned to form a perfect complementary surface to virtually any non-self epitope (2).

While the V_H_ and V_L_ domains are largely structurally symmetrical in their framework regions, the degree of symmetry is much lower in the CDR loops. Analogous CDRs of the V_L_ domain present distinct conformations when compared to those of the V_H_ (3–5). Most significant is the conformational space occupied by CDR-H3. While all other loops can be classed into distinct canonical forms (4–6), the CDR-H3 loop is highly variable and does not lend itself to structural clustering (7). This is due to the much larger sequence space explored by this loop as a result of V(D)J recombination, as the CDR-H3 loop has sequence contributions from the V, D, and J genes. The analogous CDR-L3 loop shows much lower sequence diversity due to the lack of the D segment in the light chain. Here, we use the term structural variability to refer to the diversity of CDR loop conformations found in solved crystal structures and the term flexibility to refer to the ability and likelihood of an individual CDR loop to adopt multiple different conformations.

The dissimilarities between V_H_ and V_L_ CDRs is also reflected in their contributions to the paratope, with the CDR-H3 loop generally dominating the paratope, and V_L_ CDRs making smaller contributions than their V_H_ counterparts, thereby leading to imbalanced roles of the two variable domains in antigen binding (1). Indeed, there are many cases where the V_L_ domain makes no contacts with the antigen at all (1, 8). Further, the presence of single-domain antibodies in chondrichthyes (IgNARs) (9) and camelids (V_H_Hs) (10) shows that the presence of the V_L_ domain is not necessary for antigen recognition. These observations have raised questions on the precise role of the V_L_ domain and how the presence and sequence of this region influences the V_H_ domain.

Due to the spatial proximity of the CDR-H3 loop and the V_L_ CDRs, it has long been hypothesized that one possible role of the V_L_ domain, in addition to antigen binding, might be modulating the CDR-H3 structure. This has been confirmed in specific cases, showing that the crystal structure (11) and conformational ensemble (12) of the CDR-H3 loop are indeed influenced by different pairings of V_H_ and V_L_ domains.

In this study, we show that specific attributes of the V_L_ domain influence both the structure and structural variability of CDR-H3. We first show that CDR loops are tightly packed against each other in a distance-dependent manner in antibody crystal structures and that V_L_ loops are more heavily compacted than their V_H_ counterparts. Further, we show that the CDR-H3 loop is unique in its packing, making interactions with more loops than the analogous CDR-L3 loop.

We next examine these observations in a more detailed way using a set of molecular dynamics simulations of eight antibody structures. We find that the observed packing patterns are recapitulated in simulations and that the V_L_ loops are all more rigid than their V_H_ counterparts, noting that this effect would be expected given the greater compaction of the V_L_ loops. We also show that the CDR-H3 loop explores virtually all the conformational space available to it, suggesting that the remaining CDR loops (referred to as the minor loops) act as a “cage” around the CDR-H3, constraining the number of conformations that it is able to adopt.

Using a length-independent structural similarity calculation, we show that different V_L_ genes, subtypes, and CDR canonical forms have a significant effect on the structural variability of CDR-H3, with shorter CDR-L1 and longer CDR-L3 loops increasing conformational diversity of the CDR-H3 loop and rationalise this observation in the context of our constrained CDR-H3 model. We provide a case study that shows the expected effects of V_L_ domain attributes on CDR-H3 conformation, and confirm using a structural data set that pairing the same V_H_ domain with different V_L_ sequences leads to a change in CDR-H3 conformation, with this change being correlated with the diversity of the V_L_ sequences.

## Methods

### Crystal Data Set

We mined SAbDab (13) on 09.08.2022 and filtered based on a 95% sequence identity threshold across the F_V_ domain. We further filtered this data set to exclude structures that did not have paired heavy and light chains, scF_v_s, structures not solved using X-ray crystallography and those with a resolution greater than 3Å. Lastly, we filtered the data set to remove structures with unresolved loops. Only the first F_V_ in the asymmetric unit was included in the final set. This procedure yielded 2357 non-redundant F_V_ structures.

### Region Definitions

ANARCI (14) was used to number sequences using the IMGT numbering scheme (15). We chose this numbering scheme to enable comparison across the V_H_ and V_L_ domains (CDR1: IMGT residues 27-38, CDR2: IMGT residues 56-65, CDR3: IMGT residues 105-117, Framework: IMGT residues 1-128 and not within CDR definitions).

### Crystal Radial Distribution Functions

Radial distribution functions for each loop pair were calculated by using a modified version of the mdtraj (16) implementation that does not normalise by unit cell volume. Namely, we generated histograms of pairwise heavy atom distances (up to 10Å) using a bin width of 0.1Å. We then normalised the densities by dividing the counts at each bin by the volume of the spherical shell located at that bin. This results in comparable trends but not absolute frequencies.

### Molecular Dynamics (MD) Simulations

The simulated systems were the first Fab domains in the asymmetric units of PDB entries 3gjf, 4dkf, 1e6j, 3l5x, 2cmr, 4zs7, 3l5w and 5bk1. We chose this set of antibodies as they provided a representative sample in terms of inter-loop distances (Figure 4a-j) and a range of CDR loop lengths (CDR-H1: 8-10 residues, CDR-H2: 7-8 residues, CDR-H3: 11-19 residues, CDR-L1: 5-9 residues, CDR-L2: 3 residues, CDR-L3: 8-11 residues) and sequence diversity.

#### Simulation Protocol

All systems were prepared and simulations performed using OpenMM v7.7 (17). We capped C-termini using N-methyl groups. Next, we protonated the models at a pH of 7.5, soaked them in truncated octahedral water boxes with a padding distance of 1nm, and added sodium or chloride counterions to neutralise charges and then NaCl to an ionic strength of 150 mM. We parameterised the systems using the Amber14-SB force field (18) and modelled water molecules using the TIP3P-FB model (19). Non-bonded interactions were calculated using the particle mesh Ewald method (20) using a cut-off of distance of 0.9 nm, with an error tolerance of 0.0005. Water molecules and heavy atom-hydrogen bonds were rigidified using the SETTLE (21) and SHAKE (22) algorithms, respectively. We used hydrogen mass repartitioning (23) to allow for 4 fs time steps. Simulations were run using the mixed-precision CUDA platform in OpenMM using the Middle Langevin Integrator with a friction coefficient of 1 ps^-1^ and the Monte-Carlo Barostat set to 1 atm. We equilibrated systems using a multi-step protocol: (i) energy minimisation over 10,000 steps, (ii) heating of the NVT ensembles from 100 K to 300 K over 200 ps, (iii) 200 ps simulation of the NPT ensembles at 300 K, (iii) cooling of the NVT ensembles from 300 K to 100 K over 200 ps, (iv) energy minimisation over 10,000 steps, (v) heating of the NVT ensembles from 100 K to 300 K over 200 ps, and (vi) 5 ns simulation of the NPT ensembles at 300 K.

We then initialised well-tempered metadynamics (24) using a bias height of 10 kJ/mol with width 0.3 rad and a bias factor of 10, depositing biases every 500 steps. We biased a linear combination of sine values of CDR-H3 and CDR-L3 *ψ* angles in a manner similar to (25), resulting in two collective variables. We performed triplicate simulations of 1μs duration and assessed convergence using block analysis of free energy estimates. To recover unbiased ensembles, we calculated weights (*w*) for each configuration (*s*) using the time-independent reweighting scheme (26):

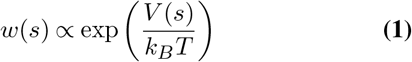

where *V* (*s*) is the bias at configuration *s* and *k*_*B*_*T* is the product of the Boltzmann constant and temperature.

#### Analysis

Analysis of MD trajectories was performed using mdtraj v1.9.6 (16) and scikit-learn v1.0.2 (27). Minimum heavy atom distances of crystal data and MD trajectories were calculated using mdtraj. MD frames were clustered using average linkage hierarchical clustering (cutoff=1.25Å) based on CDR RMSD after alignment on framework regions excluding 2 residues on either N-terminus. We chose to exclude these residues as they showed considerably higher flexibility than the rest of the framework regions. To calculate RMSD and RMSF values (defined as the time average of RMSD), we used the conformation at the beginning of the simulation as the reference.

### Dynamic Time Warping

To compare CDR-H3 loops of differing lengths, we used dynamic time warping (DTW) (28) after superposing CDR-H3 loops on anchoring residues, defined as IMGT positions 100-104 and 118-122. DTW is analogous to the Needleman-Wunsch algorithm (29), using dynamic programming to find the optimal path in a backbone atom RMSD matrix. When two loops of the same length are compared, the algorithm reduces down to a conventional RMSD calculation. When the two loops are of differing length, the optimally aligning residues are included in the comparison.

We used DTW on our crystal data set to generate a pairwise comparison matrix of all CDR-H3 loops. We then stratified the data set based on antibody subtype, F_V_ genes, and minor loop canonical forms (calculated using SCALOP (30)) and compared the average distance in the stratified data to the overall average distance. To test for significance, we used a bootstrapping procedure to randomly sample as many points from the whole data set as there were in each stratum, repeated the comparison 10,000 times and calculated a p-value based on how many times an effect of greater size was observed.

### V_H_ -V_L_ pairing analysis

To search for antibodies with identical V_H_ but dissimilar V_L_ sequences, we extracted all V_H_ sequences from SAbDab (13) on 10.12.2022 where the structure was solved using X-ray crystallography and retained only entries where F_V_ backbones were resolved completely. We clustered these antibodies on V_H_ sequences using global alignment at a clustering threshold of 1.0 using cd-hit (31). We then grouped resultant non-singleton antibodies together and clustered them again, this time using a threshold of 0.4 and clustering on the V_L_ sequences. We extracted pairs from within each subcluster if the V_L_ sequence identity was below 99%. This yielded a set of 86 antibody pairs. Every pair obtained contained at least one different residue in the V_L_ CDR loops without our method specifically selecting for pairs with CDR loop sequence identity. For each pair, we aligned the V_H_ structures on the framework regions using Biopython v1.78 (32) and calculated backbone RMSD values. Lastly, we calculated Pearson correlation coefficients using scipy v1.8.1 (33).

### Data Visualisation

We generated plots using seaborn v0.11.2 (34)and matplotlib v3.2.2 (35) and visualised protein structures using UCSF ChimeraX v1.5 (36).

## Results

### Antibody V_H_ and V_L_ CDR loops are characterised by non-equivalent inter-loop contacts in crystal structures

To analyse the structural relationships between antibody CDR loops, we began by examining a data set of antibody crystal structures. As described in the Methods section, we mined SAbDab for a set of antibodies. This yielded a set of 2357 non-redundant structures where all CDR loops were resolved. We then examined the relative packing of loops against each other by calculating radial distribution functions (RDF). For each loop, we selected heavy atoms and tallied the number of heavy atoms belonging to other loops at a given distance interval and normalised this value by dividing by the number of possible pairs. Our results are generally in line with what would be expected – loops that are close to each other in the three-dimensional antibody structure tend to make more contacts with each other and the trends are largely mirrored for the V_H_ and V_L_ loops, owing to the symmetry that marks antibody structures (Fig 1).

**Fig. 1.**
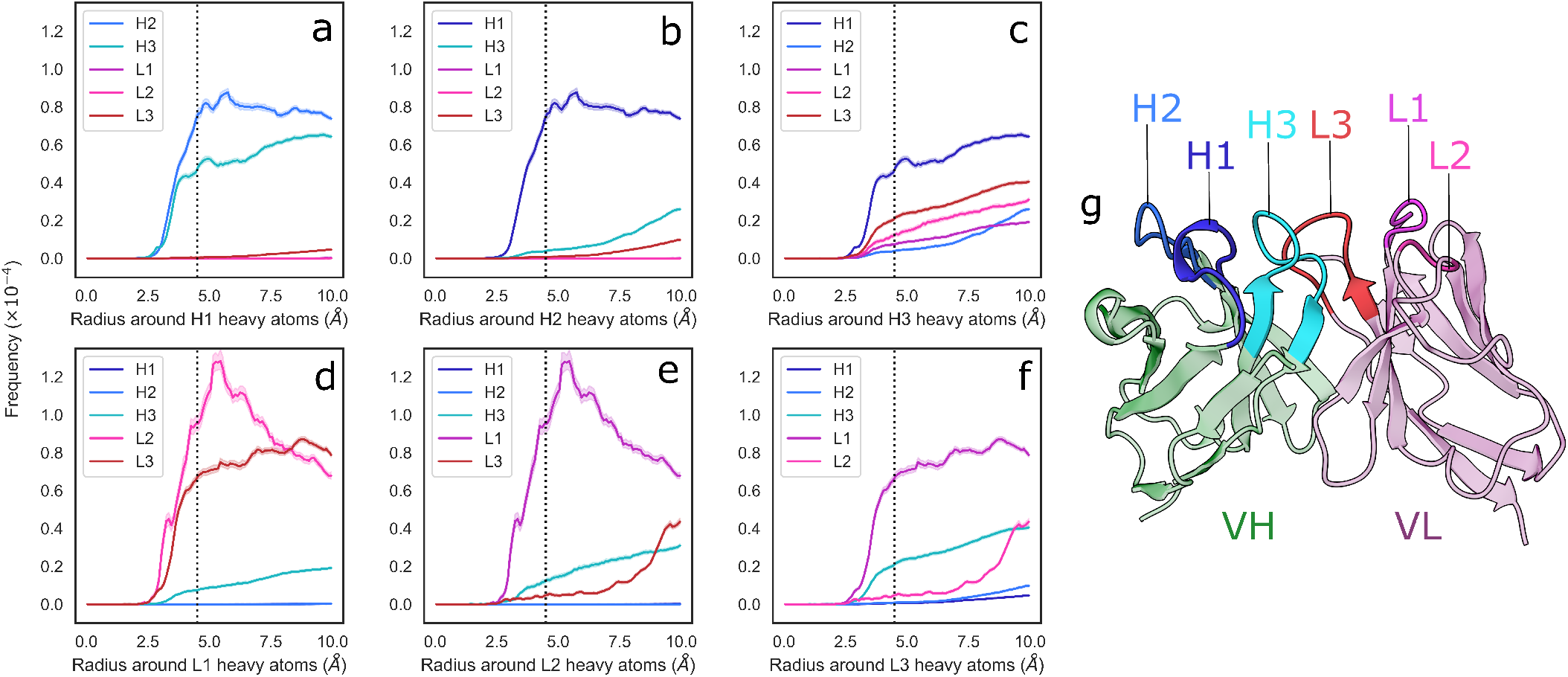
Radial distribution functions of CDR loops in the crystal data set. **(a-f)** We generated histograms for all pairwise CDR loop heavy atom distances using a bin width of 0.1Å, where **(a, b, c, d, e, f)** correspond to CDR-H1, -H2, -H3, -L1, -L2, and -L3, respectively. Each bin was then normalised by dividing by the product of the volume of a spherical shell centered at 0 and located on the edges of the bin and the number of possible pairs of heavy atoms. Shaded areas are 95% confidence intervals. **(g)** shows an example antibody structure, with CDR loops labelled according to the IMGT definition.

We found that loops sandwiched between other loops make contacts on either side. The CDR-1 loops make contacts with the CDR-2 and CDR-3 loops of the same chain, and the CDR-2 loops, owing to their positions on the edges of the VH-VL complex, only make contacts with their neighbouring CDR-1 loops (Fig 1). We did; however, find two major differences between the V_H_ and V_L_ loops.

First, that the CDR-L1/2 loops show tighter packing than the corresponding CDR-H1/2 loops. The interactions between CDR-L1 and CDR-L2 show a larger peak compared to those between CDR-H1 and CDR-H2 (Fig 1.a,d). Based on the number of residues, the CDR-L2 loop is considerably shorter (*μ* = 3.05, *σ* = 0.45) than the CDR-H2 (*μ* = 7.90, *σ* = 0.74) loop while the CDR-L1 (*μ* = 7.60, *σ* = 2.17) and CDR-H1 (*μ* = 8.19, *σ* = 0.69) loops are more comparable in their lengths. This suggests tighter packing of the CDR-L2 loop against the CDR-L1 loop than is observed with CDR-H2 and CDR-H1 based on the RDF trace, despite the shorter average length of CDR-L2.

Second, that the pattern of contacts made by the CDR-H3 loops differs significantly from that of the CDR-L3 loop. The CDR-H3 loop makes significant contacts with CDR-H1, CDR-L3, and CDR-L2 loops, with small contributions from the CDR-H1 loop (Fig 1.c), whereas the CDR-L3 loop appears to make contacts mainly with the CDR-L1 and CDR-H3 loops, with a small number of contacts made with CDR-L2 (Fig 1.f). While CDR-H3 packs less densely against CDR-H1 than CDR-L3 with CDR-L1, it packs more densely against all other loops than CDR-L3 does against its analogous loops. This is in line with the larger structural variability of CDR-H3. Thought largely to be due to their length and sequence diversity as a result of V(D)J-recombination (7), CDR-H3 loops are known to adopt multiple different conformations and can pack against minor loops in different ways. Conversely, owing to the limited conformational space afforded by the canonical forms and limited sequence space, CDR-L3 loops are more likely to pack against the same loops, in the same way (37).

We also stratified our data set on light chain subtype. We did not observe any significant differences in RDF densities in *κ*/*λ* antibodies (Fig S2), showing that the density of loop packing is the same in both V_H_ and V_L_ loops, irrespective of the light chain subtype.

Taken together, our results suggest that the V_L_ loops appear to pack more tightly against each other than the V_H_ loops and that the CDR-H3 loop is differentiated from its light chain counterpart by a much broader set of configurations with which it packs against the minor loops. We also show that the CDR loops of the V_L_ domain are more densely packed than those in the V_H_ domain, thereby likely generating a more rigid structure on one side of the F_V_.

### Conformational rearrangements in CDR-H3 do not require major conformational changes in minor loops

In order to test the observations made on crystal structures in a dynamic context, we created a set of MD simulation data. Briefly, we performed metadynamics simulations using eight antibody structures (simulation protocol details are provided in the Methods section). Each simulation was allowed to proceed for 1μs and simulations were performed in triplicates. To recover equilibrium ensembles, we re-weighted our trajectories. The simulations were biased on both the CDR-H3 and CDR-L3 loops and therefore extensive sampling of conformational space is expected for these loops only.

In the simulations the minor loops (all loops apart from CDR-H3) generally showed little flexibility. In order to quantify this, we aligned backbone atoms of each loop on three flanking residues on each side and calculated the RMSD and RMSF values relative to the starting structure. The CDR-H3 loop only appears in the starting conformation in a fraction of MD frames, with deviations ranging up to 3Å (Fig 2a). The peak that is within the 1Å cutoff used to define antibody loop conformational differences (37) is shifted further to the right for CDR-H3 compared to the major peaks found for other CDR loops (especially when compared to CDR-L3, as this loop was also biased in the simulations) (Fig 2b), suggesting that even in its native conformation, the CDR-H3 loop is more flexible and poised for conformational shifts. In line with previous studies (25, 38, 39), we therefore conclude that the CDR-H3 loop is distinctly flexible. In the context of antigen binding, where CDR-H3 is highly dominant, this might be an adaptation to allow a multitude of loop conformations in order to generate surfaces complementary to the vast range of possible epitope topologies. In our simulated systems, the least flexible CDR-H3 loop is found in non-crystal conformations in roughly 45% of frames whereas the most flexible loop immediately adopts a new conformation without revisiting the native form during the simulation (Fig 2c).

**Fig. 2.**
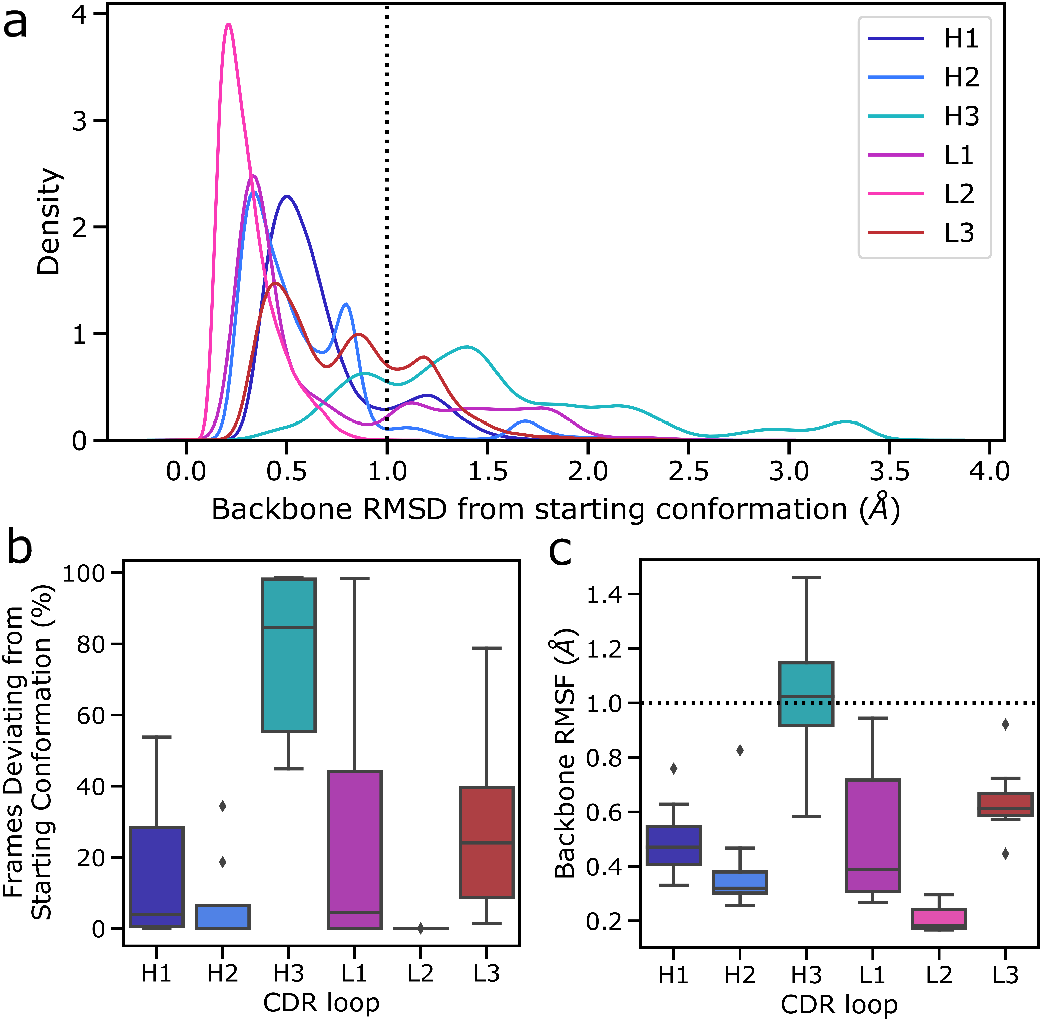
Flexibility of CDR loops in the simulation data set. All loops were indvidually aligned on their anchor residues, defined as the three residues on either side of the loop. **(a)** pooled RMSD values for each loop relative to the crystal structure (the histograms were smoothened using a kernel density estimation), **(b)** the fraction frames where loops were found in their starting conformations for each antibody, defined as being within 1Å of the crystal conformation, and **(c)** the RMSF of loops for each structure, defined as the time average of the RMSD are reported.

Conversely, the minor loops do not show similar levels of flexibility (Fig 2a). In all cases, the major peak of the distribution is within the 1Å cutoff and lies around 0.5Å. As is expected based on its length and observed conformational diversity in crystal structures, CDR-L2 is the least flexible loop, with no instances of a conformational shift being observed. The CDR-L3 loop, although biased using the same protocol as CDR-H3, is relatively rigid in its conformation. We hypothesize that this is due to a combination of factors including the length of the loop, the conformational space allowed by CDR-L3 sequences, and the tighter packing of CDR loops in the V_L_ domain. It has been shown that CDR-L3 loops tend to be shorter than CDR-H3 loops on average (37, 40). Further, a number of studies have shown that CDR-L3 loops are both less conformationally diverse and less flex-ible than CDR-H3 loops (7, 37, 41, 42). The former is presumably due to the fact that CDR-L3 loops are generated only by sequence contributions from V and J genes, whereas the D gene also contributes to CDR-H3 sequence, thereby likely increasing the conformational space by virtue of greater sequence space. The reduced flexibility compared to CDR-H3, on the other hand, is likely to be driven by a number of factors including loop length, loop sequence, and the presence or lack of stabilising lateral contacts. In line with this, a much smaller fraction of collected frames of CDR-L3 are found in non-crystal conformations (Fig 2b), though there is a significant range, with the most flexible CDR-L3 loop being in a different conformation in 80% of the trajectories. The CDR-L1 loop shows a much broader distribution in Figure 2b, especially when compared to CDR-H1, suggesting the presence of highly flexible CDR-L1 loops. However, examination of the underlying distribution shows that on average the CDR-L1 loop is found in a conformation much closer to its native pose compared to CDR-H1, as demonstrated by the shift of the major peak towards lower RMSD compared to CDR-H1 (Fig 2a).

To further quantify the observed relative flexibility of the CDR loops, we calculated RMSF values. RMSF values are the time average of RMSD relative to the starting conformation and provide a measure of flexibility. We found that in all cases, the V_L_ CDR loops show decreased flexibility relative to their V_H_ counterparts. However, our biasing scheme was highly aggressive, with biases roughly 10-fold of usual values in metadynamics simulations. This provides a high energetic incentive for conformational space exploration of the biased loops. The lack of significant conformational changes in the minor loops despite this approach suggests that conformational changes in the CDR-H3 loop are not sufficient to induce conformational changes in the minor loops. We hypothesize that the minor loops form a scaffold around the CDR-H3 loop, which then explores feasible conformations available to it based on the available space within this scaffold, as evidenced by the fact that CDR-H3 is the only loop exhibiting RMSF values beyond the 1 Å cutoff for conformational change (Fig 2c).

Our results do not indicate that the minor loops are unable to change conformation a phenomenon that has been well-documented in the literature (25, 42). Instead we discuss only the flexibility of the CDR-H3 loop when the minor loops are mainly in their crystal pose conformations. These conformations are thought to be the lowest energy shape for these loops (25, 42). Our results show that despite significant packing of the CDR-H3 loop against the minor loops, the CDR-H3 loop retains flexibility, exploring non-crystal conformations more than 40% of the time, even in the least flexible antibody in our simulation data set. Further, we show that this movement is possible without significant conformational rearrangement in the minor loops. Taken together, our results point to at least part of the conformational ensemble of CDR-H3 loops being accessible without any movement in the minor loops.

### Minor CDR loops create steric boundaries within which CDR-H3 samples conformational space

Based on our observation that CDR-H3 explores the conformational landscape in the simulation data without concurrent movement in the minor loops, we examined whether the observed packing of CDR loops in crystal structures (as shown using the RDFs in Fig 1) changes with CDR-H3 conformation. To this end, we calculated RDFs for CDR loops (Fig S1) and inter-loop minimum distances on our simulation data set (Fig 4a-j). The RDFs show that the patterns observed in the crystal data set RDFs are conserved in simulations, with CDR-H3 making a wider range of contacts compared to CDR-L3 (Fig S1c,f), and the V_L_ loops showing greater inter-loop packing than their V_H_ counterparts. This is supported by our minimum distance calculations, where in the vast majority of pairs, no significant change in the average minimum distance is observed (Fig 4a,b,c,e,f,h,j).

The RDFs for CDR-3 loops showed minor differences in interactions with CDR-2 loops. In the simulation data, we did not observe the presence of CDR-L2-CDR-H3 interaction that was found in the crystal data set (Fig 3c,d). Indeed, an examination of the minimum distance distribution for this pair (Fig 4d) shows an average increase of 2.47Å, with the majority of the distribution shifting out of the 0-4.5Å heavy atom distance range used to define interactions (43). This confirms the loss of interactions between CDR-L2 and CDR-H3 in the simulations as is shown in (Fig 4l). This is not always the case, Figure 4d exhibits a minor shoulder within the 4.5Å cutoff and Figure 4k confirms that conformations of CDR-H3 that allow interactions with CDR-L2 still exist, especially when the CDR-H3 loop is heavily oriented towards the V_L_ domain. The observation that the CDR-H3 loop generally pulls away from the CDR-L2 loop during exploration of conformational space indicates that the CDR-H3-CDR-L2 interaction might be important in the stabilisation of the crystal pose of antibodies.

**Fig. 3.**
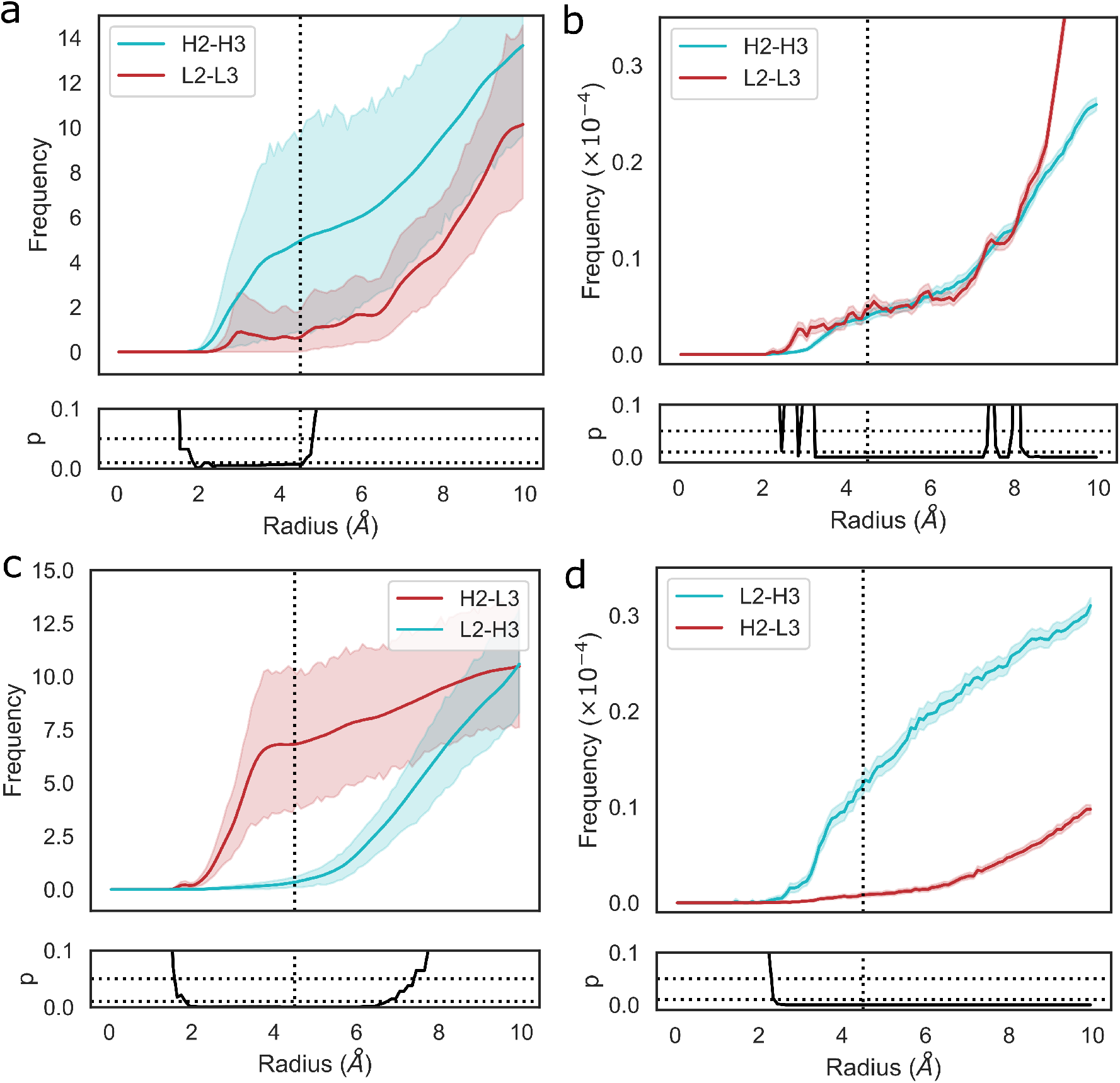
Changes in loop heavy atom packing patterns in **(a**,**c)** simulation and **(b**,**d)** crystal structure data sets. For simulation data **(a**,**c)**, we calculated RDFs using the mdtraj implementation using a bin width of 0.1Å. For crystal structure data **(b**,**d)**, we used an analogous method without the unit cell correction term. Namely, we generated histograms for all pairwise CDR loop heavy atom distances using a bin width of 0.1Å. Each bin was then normalised by dividing by the product of the volume of a spherical shell centered at 0 and located on the edges of the bin and the number of possible pairs of heavy atoms. p-values are reported based on t-tests conducted at each bin. Shaded areas correspond to 95% confidence intervals.

**Fig. 4.**
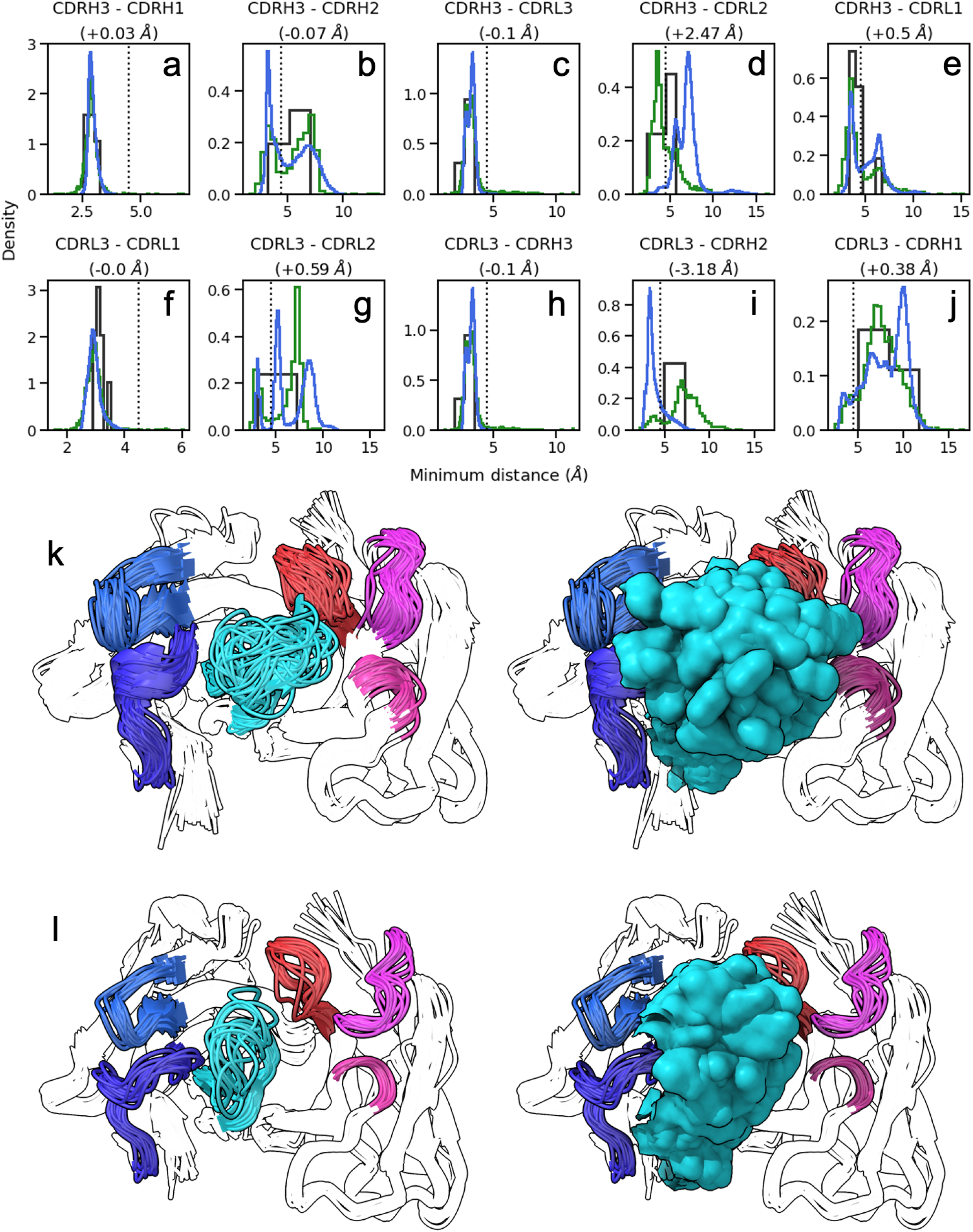
Conformations explored by CDR-H3. **(a-j)** show minimum heavy atom distances between pairs of loops. Green histograms correspond the crystal structures, blue histograms refer to simulation data, and black histograms refer to crystal structures of systems used to perform simulations. **(k**,**l)** show the conformations of CDR-H3 loops in cluster representatives extracted from simulation data clustered on CDR RMSD after alignment on the framework regions. CDR-H3 and CDR-L3 are shown in cyan and red, respectively. CDR-H1 and CDR-H2 are shown in dark and light blue, respectively. CDR-L1 and CDR-L2 are shown in magenta and pink, respectively. **(k)** shows results for 1e6j, whereas **(l)** shows results for 3l5x.

We also identified a deviation from the crystal data set in the case of the CDR-H2-CDR-L3 interaction. In this case, the opposite effect was observed, with the average minimum distance between these loops decreasing by ∼3Å on average and a concurrent shift in the simulation frame distribution to the minor peak of the crystal data distribution. This is confirmed by the appearence of density in the RDF (Fig 3c,d), showing that in simulations, CDR-L3 and CDR-H2 close the gap between them, thereby providing steric bulk in the V_H_/V_L_ interface. Based on the fact that the minor loops do not undergo any major conformational changes in our sim-ulations, this could be explained by independent changes in the framework regions of V_H_ and V_L_ domains. We found no correlation between the increase in CDR-H3-CDR-L2 and decrease in CDR-L3-CDR-H2 minimum distances.

The superimposed density maps of representatives from clustered MD trajectories show that the CDR-H3 loop samples conformations to exhaustively fill the space left by the minor loops (Fig 4k,l). This suggests that out of all possible conformations given the CDR-H3 sequence and anchor positions, the loop explores only those that are allowed within the steric mass of the minor loops. This is analogous to a “cage” formed around CDR-H3 by these loops, within which the CDR-H3 loop is able to adopt new conformations. Thus, the conformation of the other loops fine tunes the conformational landscape of CDR-H3, influencing the potential orientations of the paratope residues provided by the CDR-H3 loop. The CDR-L3 loop, being the main V_L_ loop that CDR-H3 packs against (Fig 1f, 6f) would be a major contributor to this, as the CDR-H3 and CDR-L3 loops are oriented such that they diagonally cross each other in F_V_ structures, as exemplified in Figure 6f,g. Following from the idea of the CDR-H3 loop being “caged” by the minor loops, we propose that different minor loop canonical form would lead to increases or decreases in CDR-H3 conformational diversity if a canonical form provides less or more steric bulk, thereby making the cage larger or smaller, respectively. Further, the packing between the CDR-H3 loop and minor loops provides another avenue for the modulation of CDR-H3 conformation through the presence or absence of specific interactions, in line with the observation that affinity maturation often targets residues not directly involved in contacts with the antigen, but those that make contact with other loops forming the paratope (43).

Based on our observations that CDR loops are packed against each other and CDR-H3 explores conformational space available to it within the constraints of the minor loops, we propose a model of a “caged” CDR-H3. This cage is constructed of the minor loops, with the precise orientation of these loops forming the boundaries. The CDR-H3 loop is able to undergo conformational shifts that are permitted by the exact shape and dimensions of the cage.

### Antibody subtypes, genes, and CDR canonical forms show preference for CDR-H3 conformations with different levels of diversity

Based on our “caged” CDR-H3 model, we predict that minor loops that lead to more steric bulk around the CDR-H3 loop should also lead to a reduction in conformational diversity of CDR-H3 loops. Conversely, we predict that structures with minor loop attributes that widen the physical space available to the CDR-H3 loop should lead to the broadening of conformational diversity. These predictions are in line with our results showing that CDR-H3 explores conformational space within the boundaries allowed by the minor loops in simulations. Since it is expected that the crystal pose will be part of this conformational ensemble, it follows that the steric bulk provided by minor loops should also influence the conformational diversity of CDR-H3 loops. To test these predictions, we examined whether the subtypes, genes, or minor loop canonical forms have an observable effect on CDR-H3 conformational space in crystal structures using dynamic time warping (DTW). DTW allows for the structural comparison of loops of different lengths and reduces to root mean square deviation when two loops have the same number residues. This has previously been used to effectively compare antibody CDR loops of different lengths (5) and allows us to embed the CDR-H3 conformations in our data set on a manifold in a length-independent manner. We calculated the average DTW CDR-H3 distance for the entire data set, and then stratified by subtype, gene, and canonical form before calculating average DTW CDR-H3 distances for each of these subgroups, which we then compared to baseline CDR-H3 diversity. To test for significance, we used a bootstrapping procedure.

We began by testing for differences based on light chain subtype, and found that *κ* subtypes lead to a roughly 5% reduction in the average DTW distance compared to baseline, whereas *λ* subtypes lead to a 12.5% increase, suggesting that *κ* light chains might be more likely to restrict the CDR-H3 loop. In both of these cases, we found the effect to be statistically significant with *p* < 0.05. However, we found no sigfificant differences for inter-loop packing using RDFs between *κ* and *λ* antibodies (Fig S2), suggesting that *κ* antibodies might be more likely to position minor loops in a way that restricts the CDR-H3 and that *λ* antibodies provide a larger “cage” for the CDR-H3 loop to explore conformations in. In order to examine whether specific genes drive this observation, we further stratified our data by genes. The general trend was recapitulated, but we also found that that certain genes have a higher impact on CDR-H3 conformational diversity than others. While all *λ* genes lead to an increase in average CDR-H3 diversity compared to baseline, the effect was found to be most pronounced for IGLV6, and the effect was not statistically significant in the case of IGLV1/4/5 (Fig 5c). Similarly, the majority of IGKV genes led to a reduction in CDR-H3 diversity compared to baseline, with IGKV22 being the only major exception (Fig 5b). Lastly, we examined the effect of IGHV genes. Almost all led to a modest reduction in CDR-H3 diversity, with IGHV4 as an exception (Fig 5a). This result is expected, as the IGHV gene contributes to the CDR-H3 sequence. A less diverse sequence space for the CDR-H3 loop would lead to a less diverse space of possible conformations for the loop due to residue and positionspecific preferences of backbone dihedral angles.

**Fig. 5.**
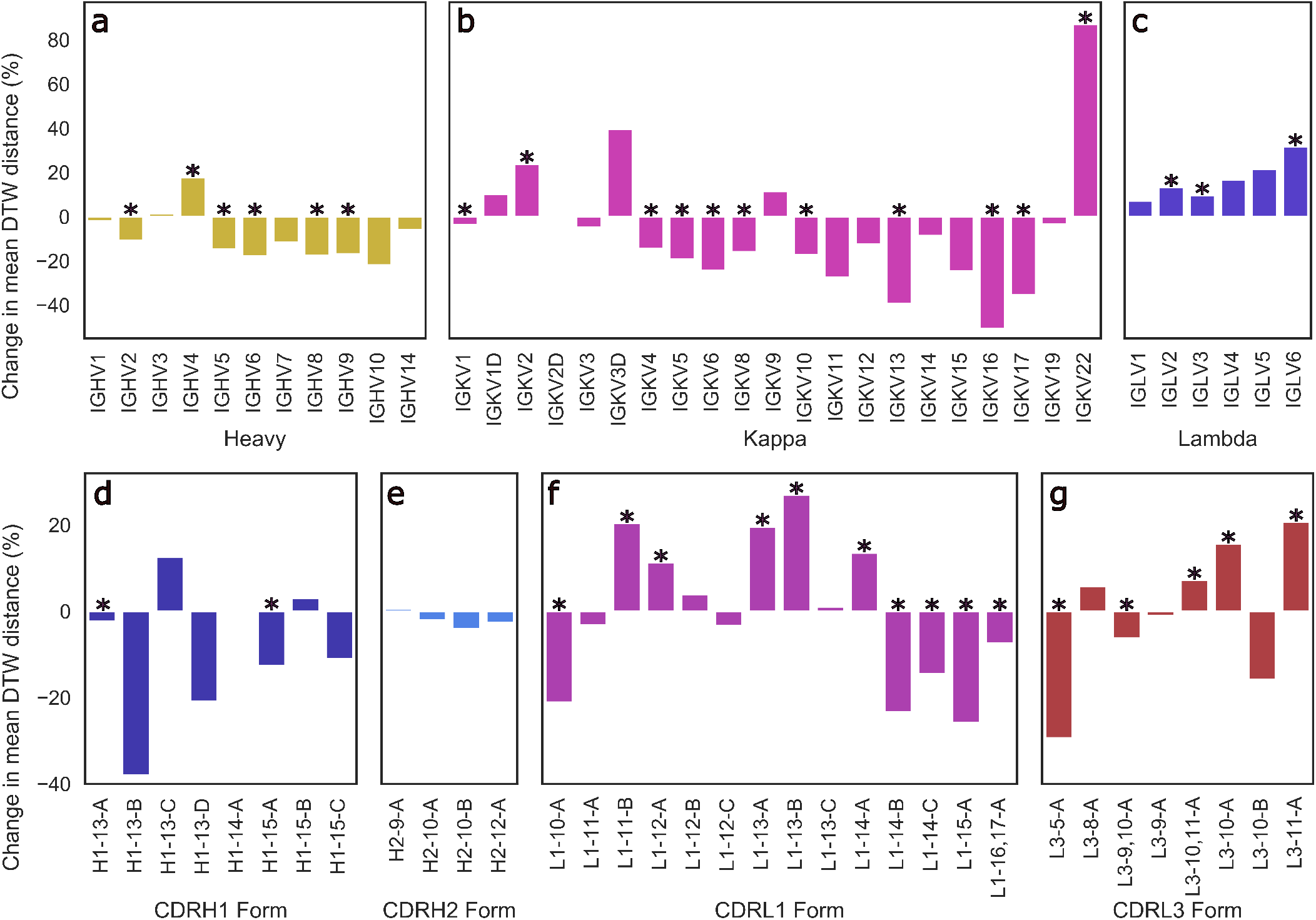
Changes in conformational diversity of CDR-H3 loops compared to baseline after stratification on **(a)** V_H_ genes, **(b**,**c)** V_L_ genes, and **(d-g)** CDR canonical forms based on dynamic time warping distances. We aligned CDR-H3 loops on anchoring residues and generated a pairwise distance matrix which was then stratified based on the examined attributes. The difference between the mean diversity of CDR-H3 loops compared to baseline (all CDR-H3 loops) is reported, with * statistical significance based on bootstrapping.

**Fig. 6.**
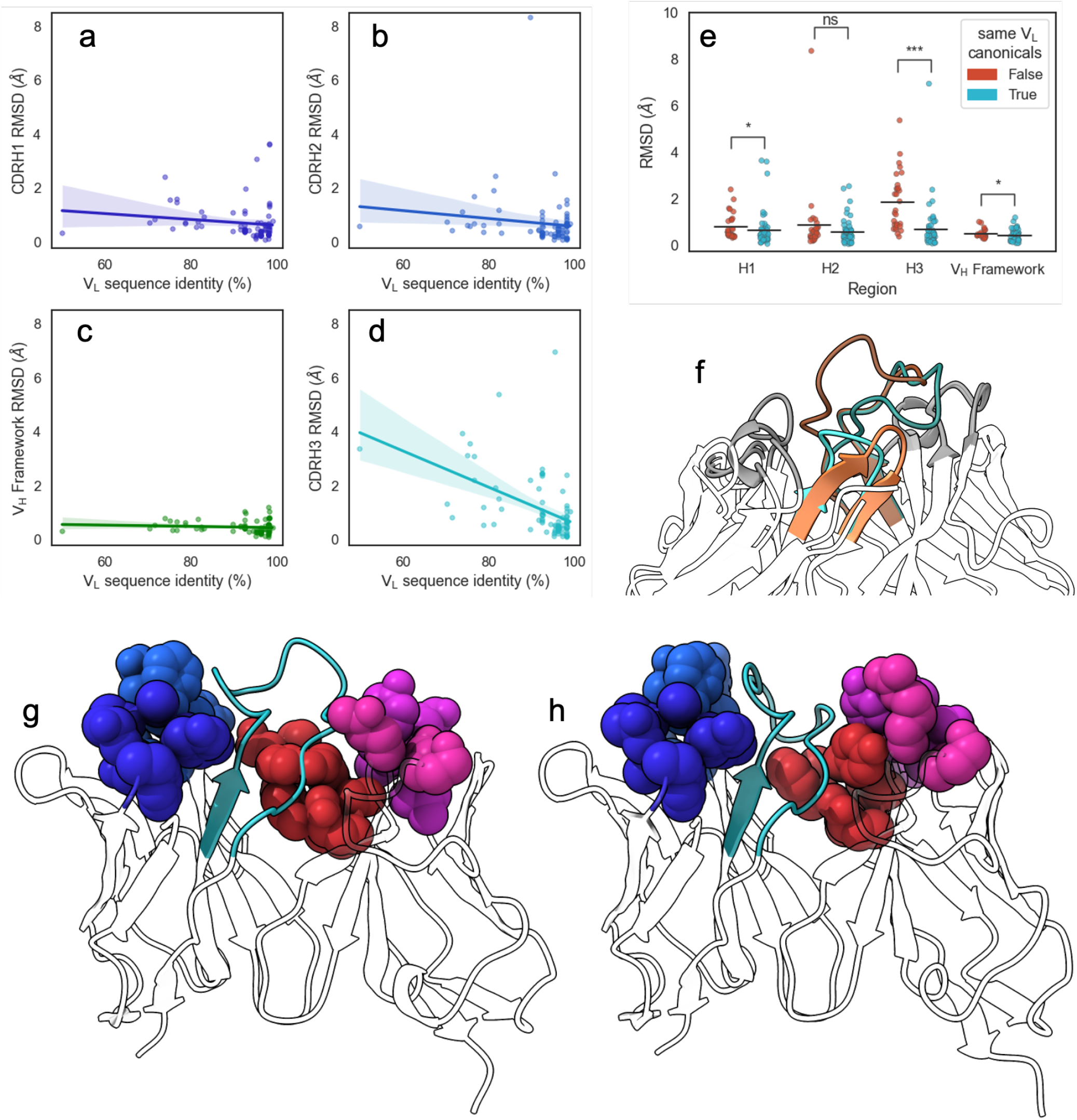
Changes in CDR-H3 conformation upon changes in V_L_ sequences. We extracted from SAbDab pairs of structures that have identical V_H_ but different V_L_ sequences and aligned structures on the V_H_ framework regions before calculating RMSD values of **(a)** CDR-H1, **(b)** CDR-H2, **(c)** V_H_ framework, and **(d)** CDR-H3 regions. Lines of best fit are reported with shaded areas corresponding to 95% confidence intervals. **(e)** shows the comparison of region RMSDs in the V_H_-identical pairs when all V_L_ loops have the same canonical forms, or when at least one loop is has a different canonical form. Statistical significance was calculated using Mann-Whitney U tests (ns: not significant, ^*^: 0.005 < *p* < 0.05, ^***^: *p* < 5 × 10^−7^) **(f)** shows the change in CDR-H3 conformation for V_H_-identical pair 5y2k (orange) and 5y2l (cyan). CDR-H3 loops are shaded using darker colors, and CDR-L3 loops are shown in lighter shades. **(g)** shows 5y2k with a shallower groove occupied by CDR-H3 due to the longer CDR-L3 and shorter CDR-L1 loop, whereas **(h)** shows 5y2l with a deeper groove occupied by CDR-H3 due to the shorter CDR-L3 and longer CDR-L1 loop. Minor loops are shown using spheres, with CDR-H3 shown in cyan.

We also examined the effect of specific canonical forms on CDR-H3 diversity. For CDR-H1 and CDR-H2, we observed that CDR-H3 diversity is limited by specific canonical forms (with H1-13-C being the single significant outlier). Since CDR-H1 and CDR-H2 canonical forms are determined by the IGHV gene, which also makes sequence contributions to the CDR-H3, we do not draw conclusions on CDR-H1-2 canonical forms on CDR-H3 conformational diversity. We also did not extend our analysis include to CDR-L2, due to its lack of diverse canonical forms.

In the case of CDR-L1 canonical forms, we observe that longer forms (14-17 residues) tend to lead to a reduction in diversity, consistent with our previous observations of loop packing. Longer CDR-L1 loops will provide more steric bulk, therefore constraining the CDR-H3 loop. Using our previously outlined framework of the minor loop ‘cage’ around the CDR-H3, this would lead to a smaller space for the CDR-H3 loop. Conversely, shorter CDR-L1 canonical forms generally lead to an increase in conformational diversity compared to baseline. For CDR-L3, we find the opposite effect: longer CDR-L3 canonical forms tend to increase the diversity of CDR-H3 conformations compared to baseline. We hypothesize that this observation is due to the relative orientation of CDR-H3 and CDR-L3. Usually in antibody crystal structures, CDR-H3 and CDR-L3 cross each other (Fig 5g, 4k,l), with CDR-H3 pointing slightly towards the V_L_ domain and CDR-L3 pointing towards the V_H_ domain. Therefore, a shorter CDR-L3 loop would enable a groove to open in the V_H_ /V_L_ (Fig 1g). As CDR-H3 makes significant contacts with CDR-L3 (shown in both our crystal data set and in our simulations), this could lead to CDR-H3 packing down towards the rest of the antibody, rather than extending further up, causing the observed decrease in CDR-H3 diversity.

These results are consistent with our model of a “caged” CDR-H3 loop, whose conformational diversity partially depends on the minor loops surrounding it, showing that CDR-H3 structural diversity is significantly modulated by V_L_ attributes as granular as CDR canonical forms.

### Identical V_H_ sequences show differences in CDR-H3 conformation when paired with different V_L_ sequences

To confirm our results from the DTW analysis of antibodies that canonical loop forms lead to a change in CDR-H3 conformational diversity, we examined a set of antibody pairs from with identical V_H_ but dissimilar V_L_ sequences, yielding a data set of 86 pairs of crystal structures, called the pairing data set (see Methods section). Briefly, we aligned each pair on the V_H_ framework region and calculated RMSD values of the V_H_ CDRs and framework region before calculating Pearson’s correlation coefficients (*r*) (Fig 6a-d). It should be noted that at the 99% V_L_ sequence identity cutoff we used (see Methods section), all pairs we identified had at least one difference in the V_L_ CDR loops. We found that the V_L_ sequence identity between these pairs was correlated significantly with CDR-H3 RMSD (*r* = −0.49, *p* < 5 × 10^−6^), but not with CDR-H1 (*r* = −0.14, *p* = 0.21), CDR-H2 (*r* = −0.13, *p* = 0.22), or V_H_ framework (*r* = −0.09, *p* = 0.40) RMSD. We also note one outlier where V_L_ sequence identity was found to be below 55%. In order to ensure that our observed correlation between sequence identity and CDR-H3 conformation is not driven by this pair, we calculated the correlation coefficient after removing it and found the correlation to still be significant (*r* = −0.46, *p* < 5 × 10^−^5).

Next, we examined the canonical forms of these pairs. We separated our data set based on whether all V_L_ canonical forms remain the same or whether at least one changes and found that changing V_L_ canonical forms lead to a greater average CDR-H3 RMSD, with an increase of 1.17 Å compared to pairs where V_L_ canonical forms remain identical. We did not observe effects of similar magnitude for CDR-H1, CDR-H2, or V_H_ framework regions, with mean RMSD increases of 0.16 Å, 0.28 Å, and 0.08 Å, respectively (Fig 6e). We found that the correlation between CDR-H3 RMSD and V_L_ sequence identity disappeared when we examined only pairs where all V_L_ canonical forms remain the same (*r*(56) = −0.11, *p* = 0.43). This suggests that even small changed in the V_L_ sequence are able to induce conformational changes in CDR-H3 without accompanying changes in the rest of the V_H_ domain. Further, we show that this is dependent on the canonical forms of V_L_ CDR loops. This supports our “caged” CDR-H3 model and provides evidence for the influence of V_L_ CDR loop steric bulk on the structure of CDR-H3 in crystal structures.

We selected the pair that showed <55% V_L_ sequence identity from this data set for further analysis: 5y2k (V_H_: chain A, V_L_: chain B) (Fig 6g) and 5y2l (V_H_: chain I, V_L_: chain J) (Fig 6h). 5y2k has one additional residue on CDR-L3 compared to 5y2l, and one fewer residue on CDR-L1. The effect of this is that the space available to CDR-H3 is much broader in 5y2k, whereas the short CDR-L3 coupled with the long CDR-L1 form the hypothesized groove that the CDR-H3 loop packs into. Indeed, when both F_V_ domains are aligned on the V_H_ domain, we observe that in the case of the shorter CDR-L3 loop, the CDR-H3 loop is folded down into the cleft and that this conformation of the loop would clash with the longer CDR-L3 loop (Fig 6f).

## Discussion

It has long been hypothesized that the conformational space of CDR-H3 loops depends not only on their sequences, but also on structural features in their vicinity. Several studies have confirmed this to be the case in specific F_V_ regions (11, 42). A question that remains is whether the presence of the V_L_ domain, or indeed specific attributes of the V_L_ domain can generally be shown to modulate CDR-H3 conformation in a predictable manner.

In this study, we show that V_L_ domains influence CDR-H3 structure by demonstrating that genetic attributes of the V_L_ domain influence both the structure and structural diversity of CDR-H3 loops. We propose a model to provide a physical explanation for this phenomenon, namely that the minor loops of the V_L_ domain can be arranged in different forms to physically restrain CDR-H3 loops, thereby modulating the conformational space that the latter are able to occupy. We argue that this is likely mediated through CDR-L loops providing steric bulk and scaffolding with which CDR-H3 can make interactions. We show that the CDR-H3 loop forms contacts with the minor loops by packing against them, and therefore hypothesize that if these minor loops were to be found in different conformations, this should be reflected in the conformation of the CDR-H3 loop. We found that in simulations where minor loop conformations do not change, the packing effects are recapitulated, in that CDR-H3 efficiently explores the conformational space that is available based on the minor loop bulk. Further, we show that inter-loop packing is tighter in the case of the V_L_, suggesting increased rigidity in the CDRs of this domain compared to the V_H_ domain. This is further supported by our observation of reduced flexibility of V_L_ loops compared to V_H_ loops.

Based on this, we show that predictable effects of minor loop attributes on CDR-H3 loop conformational diversity can be identified. We also identify the seemingly paradoxical observation that while longer CDR-L1 loops appear to reduce CDR-H3 diversity, the opposite is true for longer CDR-L3 loops and explain this observation within the context of our “caged” CDR-H3 model.

The effects observed here are one of many CDR-H3 conformation modulators, with the most significant being the specific sequence of the CDR-H3 loop. As is well-documented, the CDR-H3 loop does not lend itself to sequence-based clustering approaches suggesting that a plethora of factors are important in determining its exact structure, among which is the steric bulk provided by the minor loops. This also explains the observation that deep learning-based CDR-H3 loop modelling algorithms perform significantly better when trained using data that contains information on what is in the vicinity of the CDR-H3 loop (44). Furthermore, our observations also provide a strong case for the formulation of antibody design as a multi-objective optimisation problem where all loops must be considered together, rather than the traditional approach of modular design, where minor loops are clicked into place before subsequent modelling of CDR-H3.

In the context of B-cell immunity, our model is in line with previous studies suggesting that conformational diversity of antibodies generally decreases over the course of affinity maturation (38, 45–48). A germline combination begins with a sub-space of possible CDR-H3 conformations due to the initial F_V_ pairing (49) and over the course of the maturation process accumulates more and more mutations that home in on and stabilise the required conformation of the CDR-H3 loop for epitope complementarity.

This might then explain the observation that individual repertoires show enrichment of similar V_L_ genes upon similar antigenic insult and in functional antibodies (50), as these genes may be conducive to the necessary CDR-H3 conformational space being explored to target the antigen. However, this is only one driver of V_L_ selection, as for example in many cases the V_L_ CDR loops also make contact with the antigen, placing emphasis again on antibody design and maturation as a multi-objective problem, in both *in silico* and natural B-cell immunity contexts.

In summary, we demonstrate the role of the V_L_ CDR loops as modulators of the CDR-H3 loop through modifications of the conformational subspace of the loop. Further research into this role and its examination in dynamical studies of antibodies where minor loop flexibility is also explored will allow us to more accurately understand the interconnected nature of CDR loop conformations. This will be important to fully understand the effect of this on rational antibody design.

## AUTHOR CONTRIBUTIONS

B.G. and C.M.D. conceived the project and designed the study. B.G. performed experiments. B.G. and C.M.D. wrote the manuscript. C.M.D supervised the project.

