## Supplementary Figures for "Specific attributes of the V_L_ domain influence both the structure and structural variability of CDR-H3 through steric effects"

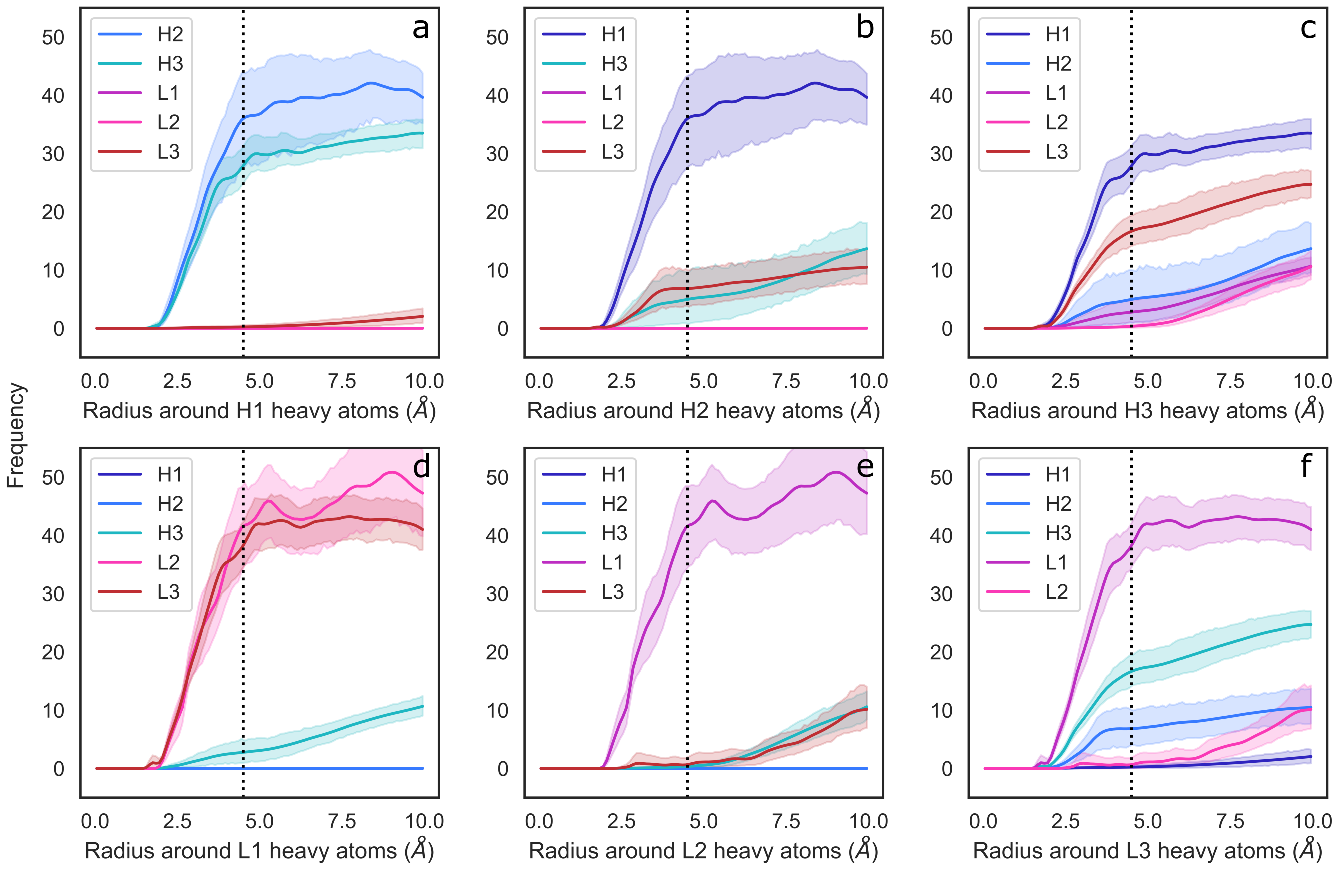


**Fig. S1.** Radial distribution functions of CDR loops in the molecular dynamics data set, calculated RDFs using the mdtraj implementation using a bin width of 0.1Å. Shaded areas correspond to 95% confidence intervals.

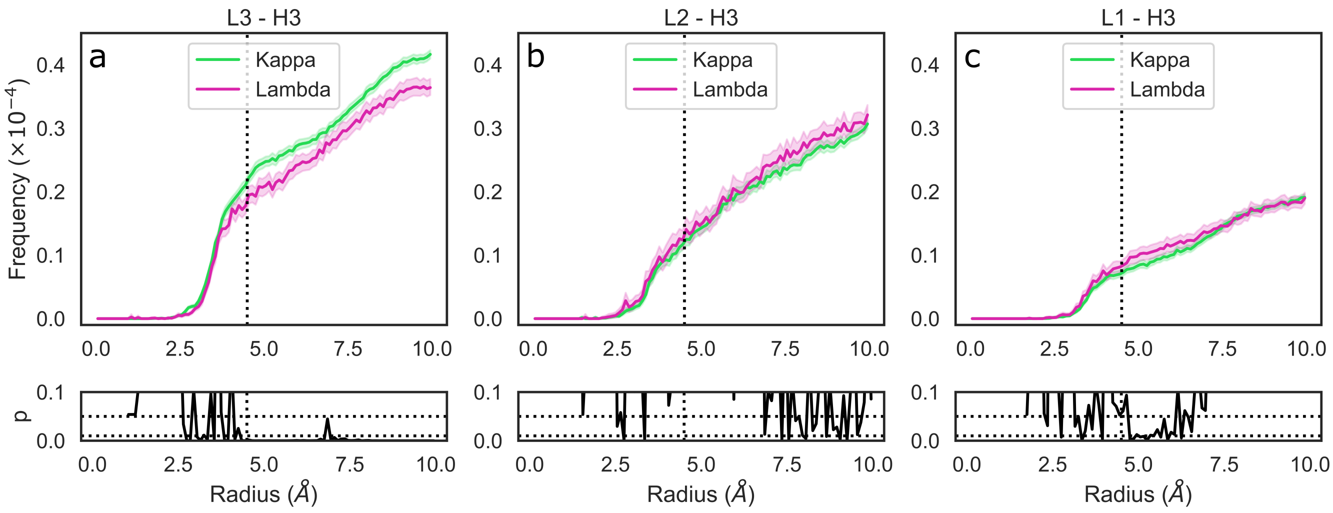


**Fig. S2.** Radial distribution functions of CDR loops in the crystal data set separated by light chain type. e generated histograms for all pairwise CDR loop heavy atom distances using a bin width of 0.1 Å. Each bin was then normalised by dividing by the product of the volume of a spherical shell centered at 0 and located on the edges of the bin and the number of possible pairs of heavy atoms. Shaded areas are 95% confidence intervals.
